# Engineering sensitivity and specificity of AraC-based biosensors responsive to triacetic acid lactone and orsellinic acid

**DOI:** 10.1101/2020.09.15.299107

**Authors:** Zhiqing Wang, Aarti Doshi, Ratul Chowdhury, Yixi Wang, Costas D. Maranas, Patrick C. Cirino

**Affiliations:** Dept. of Chemical and Biomolecular Engineering, University of Houston, Houston, TX; Dept. of Biology and Biochemistry, University of Houston, Houston, TX; Dept. of Chemical & Biomedical Engineering, Penn State University, University Park, PA

**Keywords:** Synthetic biology, Protein engineering, Polyketides, High throughput screening, Molecular reporters, IPRO

## Abstract

We previously described the design of triacetic acid lactone (TAL) biosensor “AraC-TAL1”, based on the AraC regulatory protein. While useful as a tool to screen for enhanced TAL biosynthesis, this variant shows elevated background (leaky) expression, poor sensitivity, and relaxed inducer specificity, including responsiveness to orsellinic acid (OA). More sensitive biosensors specific to either TAL or OA can aid in the study and engineering of polyketide synthases that produce these and similar compounds. In this work, we employed a TetA-based dual-selection to isolate new TAL-responsive AraC variants showing reduced background expression and improved TAL sensitivity. To improve TAL specificity, OA was included as a “decoy” ligand during negative selection, resulting in isolation of a TAL biosensor that is inhibited by OA. Finally, to engineer OA-specific AraC variants, the IPRO computational framework was employed, followed by two rounds of directed evolution, resulting in a biosensor with 24-fold improved OA/TAL specificity, relative to AraC-TAL1.

## Introduction

Regulatory protein-based biosensors that respond to small molecule ligands have been used to engineer and optimize enzymes and biosynthetic pathways (Rogers, Taylor and Church 2016, Dietrich, McKee and Keasling 2010, Zhang and Keasling 2011, Wang and Cirino 2016). Two major applications for biosensors in metabolic engineering are: (1) to facilitate high-throughput screening/selection of large enzyme/pathway libraries; and (2) dynamic regulation to balance cell growth and biochemical production. In both scenarios, the sensor to be employed must be equipped with the relevant ligand sensitivity and specificity. However, engineered regulatory proteins often display lowered sensitivity toward the non-native ligands and relaxed specificity, as compared to their wild-type counterparts (Collins, Leadbetter and Arnold 2006, Reed, Blazeck and Alper 2012, Tang et al. 2013, Chen et al. 2015, Lönneborg, Varga and Brzezinski 2012, Taylor et al. 2016). Furthermore, many engineered regulatory proteins display elevated background (leaky) expression levels (Lönneborg et al. 2012, Chen et al. 2015, de los Santos et al. 2016); for some applications, high background expression can be detrimental (Loew et al. 2010, Lachmann et al. 2015).

Regulatory protein AraC was previously engineered to respond to non-native ligands, and those variants facilitated high-throughput screening of enzymes and pathways (Tang et al. 2013, Tang and Cirino 2011, Qian, Li and Cirino 2019). One AraC variant, AraC-TAL1, responds to triacetic acid lactone (TAL). AraC-TAL1 was used in blue/white screening of libraries of the type III polyketide synthase (PKS), 2-pyrone synthase (Tang et al. 2013), as well as screening of host genome libraries (gene deletions and overexpression) (Li et al. 2018), for enhanced TAL production by engineered *E. coli*. This TAL sensor displays elevated background expression as compared to wild-type AraC, as well as relaxed substrate specificity and relatively low sensitivity to TAL (half-maximum dose response occurs at 4 mM TAL concentration). More interesting applications of a TAL biosensor, or a biosensor that responds to similar, minimal polyketides such as orsellinic acid (OA), lie in the identification of novel PKS variants whose poor functional expression and/or low catalytic activity demand significantly greater sensitivity and/or specificity than AraC-TAL1 (Yeom et al. 2018). A similar constraint was recently described by Thompson *et al*., for the case of engineering biosynthesis of the nylon precursor caprolactam (Thompson et al. 2020). A highly sensitive caprolactam biosensor identified from *Pseudomonas putida* (K_d_ ≈ 5 μM) proved useful for detecting minute caprolactam levels, enabling rapid and accurate screening of novel production pathways (Thompson et al. 2020).

In the present study, we first sought variants of AraC-TAL1 showing reduced background expression and higher TAL sensitivity, using a TetA-based dual selection to rapidly enrich a library of ligand binding domain variants. Seven variants of AraC-TAL1 having improved sensitivity and/or lowered background expression (comparable to that of wild-type AraC) were isolated using this selection platform. All seven AraC-TAL1 variants show various degrees of relaxed inducer specificity, including response to OA. A subsequent round of selection in which OA was included as a decoy ligand resulted in isolation of variant AraC-TAL^+^/OA^-^ (carrying nine total amino acid substitutions relative to wild-type AraC), in which OA competitively inhibits the TAL-induced response.

OA is a common tetraketide product of type I PKS systems (Sanchez et al. 2010), and an OA-producing type III PKS from *Rhododendron dauricum* has been described (Taura et al. 2016). Importantly, TAL is a common “derailment” triketide byproduct during OA biosynthesis (Taura et al. 2016). An OA sensor useful for engineering OA biosynthesis, or for studying/engineering chain elongation vs termination during tetraketide biosynthesis, must therefore show specificity toward OA and not TAL. We accordingly sought variants with higher specificity toward OA, using computational protein design followed by directed evolution. Variant AraC-OA8, with 12 total amino acid substitutions relative to wild-type AraC, shows a 24-fold increase in the OA specificity as compared to AraC-TAL1 and retains low background expression, comparable to that of wild-type AraC. Collectively, the new AraC variants described have potential utility as biosensors for engineering PKS specificity and improving OA biosynthesis, as well as transcriptional regulators in related engineered biosynthesis pathways.

## Materials and Methods

### Plasmid Construction

Plasmid pPCC1322 (map and sequence in Supplementary Materials) was constructed from pFG29-TAL (Frei et al. 2016) as follows: the f1 origin, together with ∼1 kb non-coding sequence flanking the f1 origin, was removed from pFG29-TAL, and the ribosome binding site (RBS) region of AraC-TAL1 was modified to achieve a lower translation initiation rate for AraC-TAL1 or its variants. Library enrichment with *tetA-*dual selection was carried out with plasmids pPCC1340 and pPCC1342. Plasmid pPCC1340 was constructed by swapping *gfp* of pPCC1322 with *tetA. TetA* was amplified from PC05 chromosome (Khankal et al. 2009). Plasmid pPCC1342 was constructed by replacing the RBS of *tetA* with a stronger RBS (designed with RBS Calculator). For comparison, the TetA translation initiation rates, calculated using the RBS Calculator (Espah Borujeni, Channarasappa and Salis 2014, Espah Borujeni and Salis 2016, Espah Borujeni et al. 2017, Salis, Mirsky and Voigt 2009) (https://salislab.net/software/), are 20607 au with plasmid pPCC1340 and 571456 au with pPCC1342.

T4 ligase and all restriction enzymes were purchased from New England Biolabs. T4 ligase was used for ligation reactions. NEBuilder® HiFi DNA Assembly Master Mix was used for Gibson Assembly (Gibson et al. 2009). High-fidelity PCR in this work was performed using Phusion® High-Fidelity DNA Polymerase or Q5® High-Fidelity DNA Polymerase. PCR conditions followed NEB Tm Calculator (https://tmcalculator.neb.com/) and vendor’s instructions for the polymerases. Zymoclean^™^ Gel DNA Recovery Kit was used for gel purification of DNA fragments.

### Library Construction

Random mutagenesis libraries were generated by amplifying the ligand binding domain (LBD) of the respective parent AraC variant using the Agilent GeneMorph® II Random Mutagenesis Kit. For TetA selections, the library insert was ligated into pPCC1342, in place of the gene encoding AraC-TAL1. For libraries with AraC-OA6 and AraC-OA7 as parents (GFP fluorescence-based screening in microtiter plates), the library inserts were ligated into pPCC1322 vector. Library error rates were determined by sequencing ten random library clones: error rates were ∼4.5 mutations/kb for TetA selection libraries, and ∼4.2 mutations/kb for GFP screening libraries.

Combinatorial assembly of mutations identified in variants AraC-TAL12 to AraC-TAL17 was performed by assembly of purified PCR fragments amplified from each mutant (including parent AraC- TAL1). Primers were designed such that each fragment covers one mutation. For two mutations that are in proximity, fragments were amplified by PCR such that the fragments include all four possible combinations of the two mutations. The fragments were then assembled with Gibson Assembly into pPCC1322 vector for fluorescence-based microtiter plate screening. Sequencing of randomly picked clones confirmed correct library assembly. A nearly identical assembly approach was used to construct the library representing all 32 combinations of the five single- or double-amino acid substitutions returned by IPRO (those in AraC-OA1 through AraC-OA5).

### Iterative Protein Redesign and Optimization to Isolate OA-specific AraC-based Biosensors

Using the modeled structure of AraC-TAL14 as the starting point for computational design, OA-specific AraC variants were obtained using an iterative protein redesign and optimization approach (IPRO (Pantazes et al. 2015)), with unrestricted choice of substituted amino acid type for positions Pro8, Pro11, Thr24, Pro25, Gly30, Leu72, His80, Tyr82, Arg89, His93, Gly135, and Asn139. These residue positions include those that have previously been targeted for saturation mutagenesis (Tang, Fazelinia and Cirino 2008), in addition to residue positions that appeared in new TAL-specific AraC variants described in this work. The protein redesigns were driven by the primary objective of enhancing OA interaction (sequence redesign step) and secondary objective of eliminating TAL interaction (only side chain repacking without introducing further amino acid changes). Note that ligand binding in AraC does not necessarily induce the requisite conformational changes to activate gene expression. Hence, ligand-interacting AraC variants with no induced GFP expression are expected to be encountered during experimental characterization of the IPRO-returned designs. After sampling several design trajectories that progressively incorporate amino acid substitutions that improve binding to OA without destabilizing the variant structure (by more than 25% of the AraC-TAL14), the top five designs with highest OA/TAL interaction energy ratios and with single or double amino acid substitutions, were chosen for experimental characterization.

### Fluorescence Assays for Measuring GFP Expression

GFP expression under the control of AraC-based biosensors was measured using a microtiter plate-based fluorescence assay. For characterizing individual variants and constructing dose response curves: single colonies of HF19 (Tang et al. 2008) transformants were inoculated into 500 µL LB supplemented with 50 µg/mL apramycin, in 96-deep-well-plates. After 6-12 hours of growth at 37 °C 900 RPM on a shaking platform, the cultures were then diluted into fresh LB containing 50 µg/mL apramycin and 100 µM IPTG, with or without inducer ligand(s) of interest at final concentrations as indicated in the presented results. The subcultures were grown for 4-6 hours at 37 °C 900 RPM in 96-well deep well plates. The cells were next pelleted and washed with PBS buffer before measurements. Fluorescence was measured on SpectraMax® Gemini^™^ EM Microplate Spectrofluorometer from Molecular Devices®. Optical Density at 595 nm (OD_595_) was measured on BMG Labtech NOVOstar Microplate reader. Fluorescence intensity was normalized by OD_595_. Fold-induced GFP expression was calculated by dividing the normalized fluorescence intensity in presence of the inducer by the normalized fluorescence intensity in absence of the inducer.

For microtiter plate-based fluorescence screening of AraC libraries, single colonies of fresh HF19 transformants (450-500 colonies from each library) were inoculated into 500 µL LB supplemented with 50 µg/mL apramycin, in 96-deep-well-plates. HF19 cells transformed with pPCC1322 containing wild-type AraC or AraC-TAL14 in place of AraC-TAL1 were used as controls on each screening plate. The library cultures were then handled as described above for the case of individual clones. For the small combinatorial library comprising IPRO-predicted substitutions, three sets of subcultures were prepared for screening: with no inducer, 0.5 mM OA, or 3 mM TAL. For the random mutagenesis library, subcultures were prepared with no inducer, 1 mM OA, or 3 mM TAL. Top clones from the combinatorial library were defined as those which had the highest OA specificity ratio (ratio of fold-induced GFP with OA to the fold-induced GFP with TAL), while the best clones from the random mutagenesis library were those with the highest fold-induced GFP with OA. All data points reported represent the average values of at least two independent replicates.

### TetA Dual-selection Conditions

#### Enriching AraC-TAL1 variants with reduced background expression and higher sensitivity

HF19 competent cells were transformed with the library plasmid pool and the transformants were inoculated into 500 mL LB supplemented with 50 µg/mL apramycin, 100 µM IPTG, and 1 mM NiCl_2_. The negative selection was performed with vector pPCC1342 (strong *tetA* RBS), starting with ∼10^8^ transformants and harvested when OD_595_ was above 1 (total ∼2.5×10^11^ CFU). After each round of negative selection, the library was harvested, and the plasmids were extracted. The library insert was then re-cloned into the pPCC1340 backbone (weak *tetA* RBS) for positive selection on LB-Agar plates in the presence of 50 µg/mL apramycin, 100 µM IPTG, 1% glycerol, 50 mM TES (pH 7), 8 µg/mg Tc and 0.5 mM TAL. Library transformants were plated directly on positive selection plates and cells were harvested by scraping after ∼13 hours of incubation at 37 °C for plasmid extraction. The harvested library then was re-cloned back to pPCC1342 vector for the next round of negative selection. After three rounds of dual-selection, the enriched library was cloned to pPCC1322 backbone for an end-point screening with fluorescence assay in 96-well deep-well plates. 94 colonies of HF19 transformants with the enriched library were assayed in the fluorescence assay for their leakiness and responses to TAL.

#### Enriching AraC-TAL1 variants that do not respond to orsellinic acid

Here, both negative and positive selections were carried out using pPCC1340 (weak *tetA* RBS). Negative selection was carried out in LB supplemented with 50 µg/mL apramycin, 100 µM IPTG, 0.5 mM NiCl_2_ and 2 mM orsellinic acid, starting with ∼10^8^ transformants and harvested when OD_595_ was above 1 (total ∼2.5×10^11^ CFU). The plasmids were then extracted for re-transformation for positive selection (re- transformation was performed to eliminate any potential mutations appearing in the host genome). Positive selection was carried out on LB-Agar plates as described above. After three rounds of dual-selection, the enriched library was cloned to pPCC1322 backbone for an end-point screening with fluorescence assay in 96-well deep-well plates. 94 colonies of HF19 transformants with the enriched library were assayed in the fluorescence assay for their responses to 1 mM TAL and 1 mM OA.

## Results

### TAL Sensors with Improved Sensitivity and Reduced Background Expression

The *tetA*-encoded class C tetracycline resistance protein (TetA) is a tetracycline/H^+^ antiporter that confers simultaneous tetracycline (Tc) resistance and nickel (Ni^2+^) sensitivity to *E. coli* (Stavropoulos and Strathdee 2000). These combined features have enabled development of dual selection systems in *E. coli* based only on *tetA* expression, for engineering riboswitches (Nomura and Yokobayashi 2007, Muranaka et al. 2009) and genome editing (Ryu et al. 2017). Attracted to its simplicity for library screening, we optimized the *tetA* dual selection system to isolate AraC-based biosensors showing improved TAL sensitivity and reduced background expression, as compared to AraC-TAL1 (Figure 1A). The gene encoding the AraC-TAL1 ligand binding domain (LBD) was amplified using error-prone PCR, and the random mutation library was expressed in *E. coli* strain HF19. Expression of *tetA* was placed under control of promoter P_BAD_, regulated by an AraC-TAL1 variant expressed from the same plasmid. AraC-TAL1 variants allowing leaky expression of *tetA* in absence of TAL (and in presence of Ni^2+^) are eliminated by virtue of their Ni^2+^ sensitivity during negative selection. Meanwhile inclusion of Tc (8 μg/mg) allows for positive selection of AraC-TAL1 variants induced by a relatively low concentration of TAL (0.5 mM). Iterative rounds of negative selection enriched AraC-TAL1 variants having low background in absence of TAL, with positive rounds added after each negative selection step, to enrich only variants responsive to TAL.

**Figure 1.**
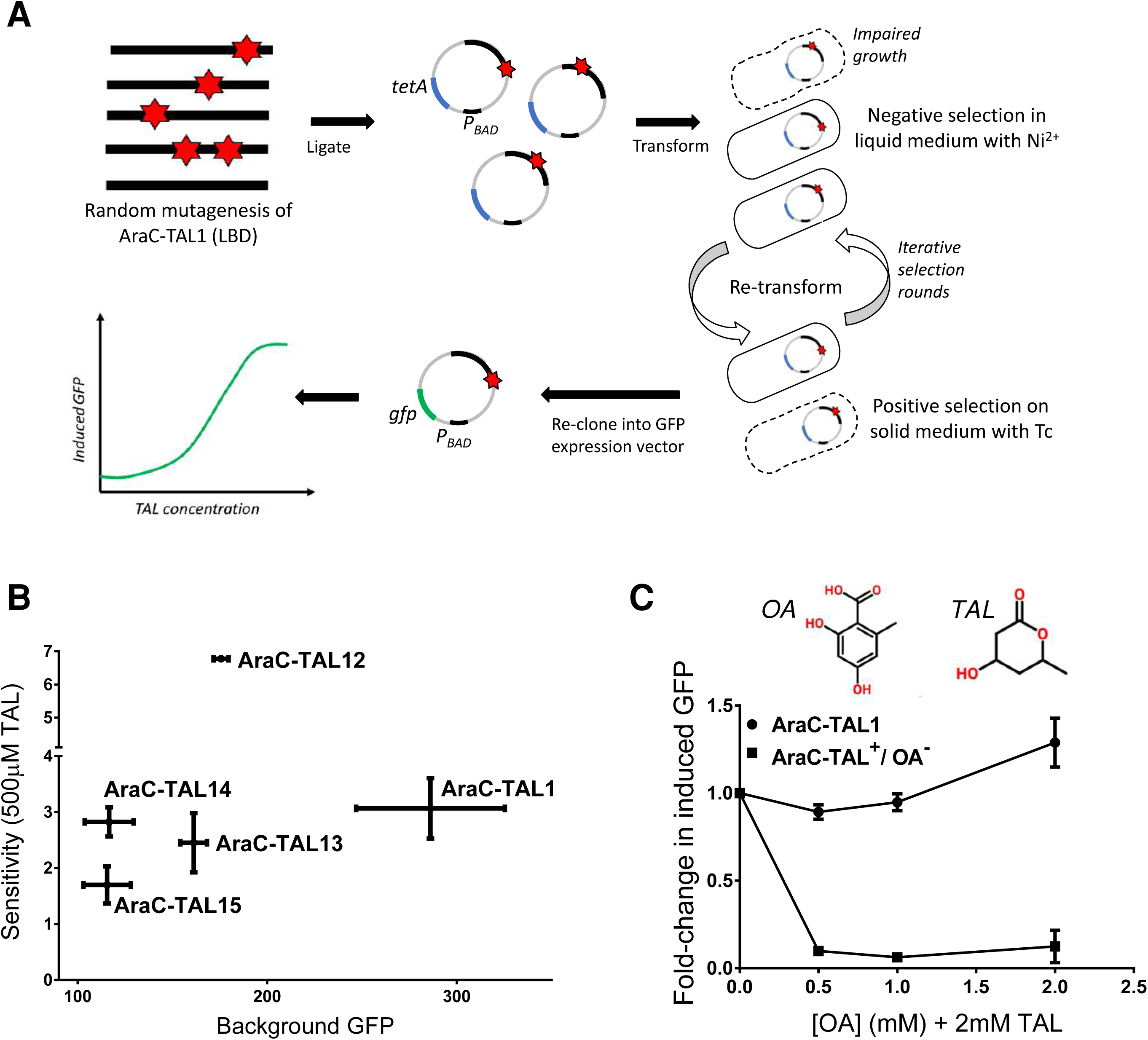
AraC-based TAL sensors with improved sensitivity and reduced background expression. A) Iterative rounds of negative selection (in presence of Ni^2+^ and absence of TAL) in liquid medium and positive selection (in presence of Tc and TAL) on solid medium to isolate AraC-TAL1 variants with higher sensitivity and lower background expression as compared to AraC-TAL1. The selected mutants were cloned into a P_BAD_-*gfp* expression vector for end-point screening and fluorescence-based dose response characterizations. B) TAL sensors with improved sensitivity and/or reduced background expression, relative to AraC-TAL1. Sensitivity is defined as the induced GFP expression response when cells were grown in media supplemented with 500 μM TAL. Background expression for wild-type AraC is 89 ± 3. C) OA inhibits TAL-induced GFP expression in variant AraC-TAL^+^/OA^-^. All fluorescence measurements were normalized by the measured cell density (RFU/OD595). Background GPF represents the fluorescence measurements (RFU/OD595) in the absence of inducer. Induced GFP expression indicates the normalized fluorescence value in presence of the inducer divided by the normalized fluorescence value in absence of the inducer. Data points are the average of two values, and error bars represent the range.

Following three rounds of dual selection, we identified six AraC-TAL1 variants (named AraC- TAL12 to AraC-TAL17) having higher sensitivity and lower background expression than the AraC-TAL1 parent. Sequencing of the selected mutants identified single amino acid substitutions in the AraC-TAL1 LBD (Table 1). These six AraC-TAL1 variants were further characterized using a P_BAD_-*gfp* expression vector, for TAL-dependent fluorescence measurements (results summarized in Figure 1B; dose response curves shown in Figure S1). AraC-TAL12 shows >2-fold greater induced GFP expression with 0.5 mM TAL, as compared to the AraC-TAL1 parent (6.8-fold vs. 3.1-fold). Meanwhile AraC-TAL14 and AraC-TAL15 show the tightest repression in the absence of TAL, with background GFP expression levels comparable to that of wild-type AraC (Figure 1B).

**Table 1.**
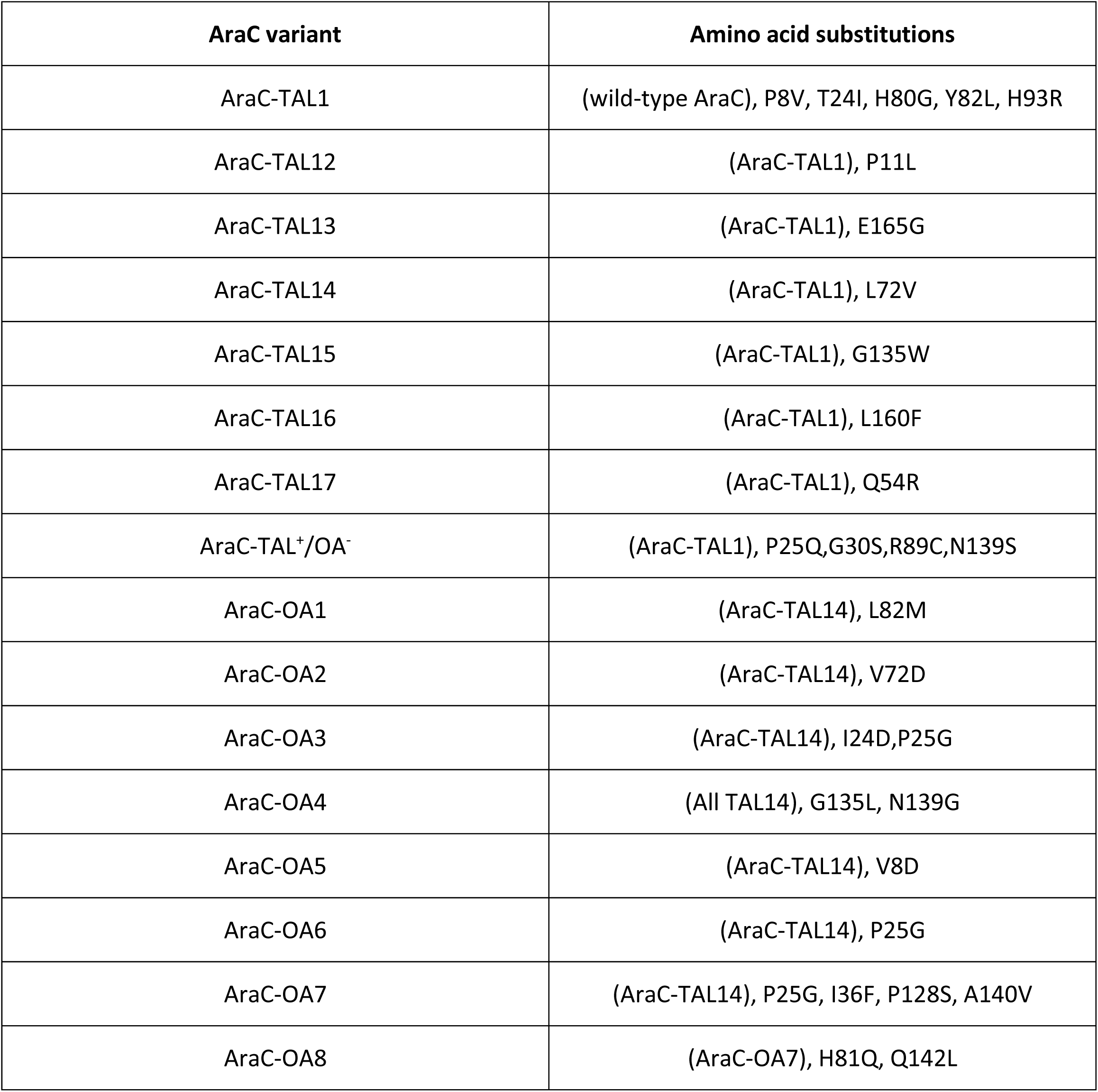
Amino acid substitutions in AraC-based biosensors described in this work.

A small library was next constructed to test whether any combinations of the amino acid substitutions in variants AraC-TAL12 to AraC-TAL17 would further improve sensitivity or reduce background expression. Details of library construction and screening are provided in Materials and Methods. With only 64 possible combinations, the P_BAD_-*gfp* expression vector was used for library cloning, and 280 total transformants were screened by microtiter-plate-based fluorescence measurement in the presence vs absence of 1 mM TAL. The individual amino acid substitutions appear to be largely non-additive since all clones showed TAL response and background GFP expression levels similar to or worse than those of the parent clones. Results from screening and characterization of one variant (named AraC- TAL18) carrying substitutions from AraC-TAL12 (P11L) and AraC-TAL17 (Q54R) are presented in the Supplementary Material (Table S1).

### A TAL Sensor with Improved Specificity

Our previous studies show that our AraC-based TAL sensors also respond to one or more compounds that are structurally similar to TAL, such as 4-hydroxy-3,6-dimethyl TAL (4H36M TAL), and 2-hydroxybenzoic acid (salicylic acid) (Frei et al. 2016, Frei, Qian and Cirino 2018). We find similar ligand promiscuity for these new TAL sensors, with results summarized in Table 2 (response curves are given in Figure S2 and Figure S3). Variants AraC-TAL12 through AraC-TAL17 all respond to 2-hydroxybenzoic acid (2OHBA or salicylic acid), 4H36M TAL, and 2,4-dihydroxy-6-methylbenzoic acid (orsellinic acid), to various extents, as does AraC-TAL1. Meanwhile, none show response to 4-hydroxybenzoic acid (4OHBA), L-arabinose, or phloroglucinol. There is no apparent correlation between response to TAL and response to other inducers tested. As examples, AraC-TAL16 shows a response to 2 mM TAL that is ∼2-fold higher than that of AraC- TAL14 (Figure S1), but its response to 2 mM orsellinic acid is ∼2-fold lower than that of AraC-TAL14 (Figure S2B). Similarly, AraC-TAL12 responds to 2 mM TAL 3.3-fold stronger than AraC-TAL1 (Figure S1), but response to 2 mM orsellinic acid is similar to that of AraC-TAL1 (Figure S2B). These observations demonstrate how single amino acid substitutions can alter not only sensitivity but also specificity of the ligand response.

**Table 2.**
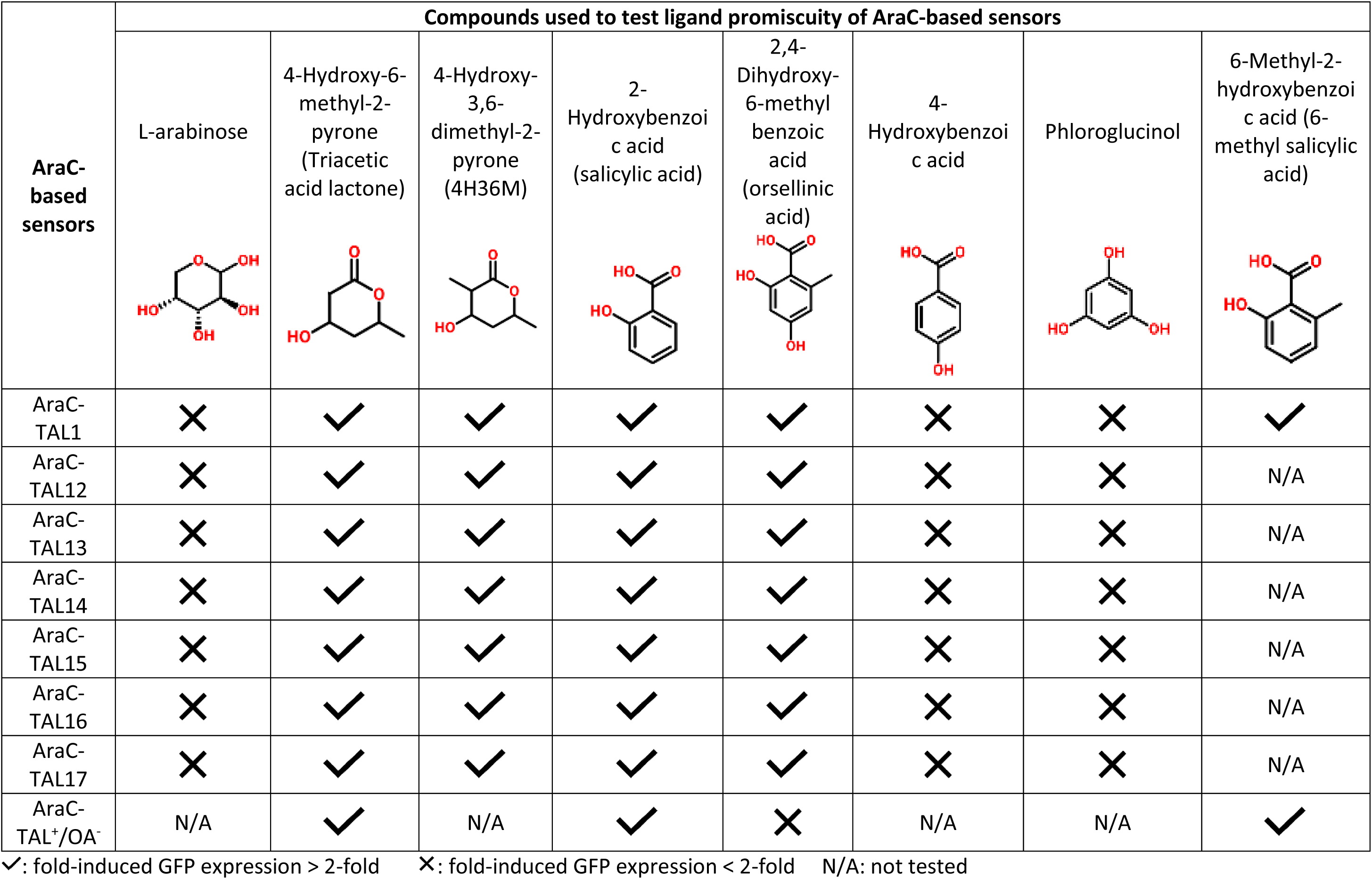
Analyzing the ligand promiscuity of the AraC-TAL1 variants using several compounds.

| AraC-based sensors | Compounds used to test ligand promiscuity of AraC-based sensors |  |  |  |  |  |  |  |
| --- | --- | --- | --- | --- | --- | --- | --- | --- |
|  | L-arabinose | 4-Hydroxy-6-methyl-2-pyrone (Triacetic acid lactone) | 4-Hydroxy-3,6-dimethyl-2-pyrone (4H36M) | 2-Hydroxybenzoic acid (salicylic acid) | 2,4-Dihydroxy-6-methylbenzoic acid (orsellinic acid) | 4-Hydroxybenzoic acid | Phloroglucinol | 6-Methyl-2-hydroxybenzoic acid (6-methyl salicylic acid) |
| AraC-TAL1 | ✗ | ✓ | ✓ | ✓ | ✓ | ✗ | ✗ | ✓ |
| AraC-TAL12 | ✗ | ✓ | ✓ | ✓ | ✓ | ✗ | ✗ | N/A |
| AraC-TAL13 | ✗ | ✓ | ✓ | ✓ | ✓ | ✗ | ✗ | N/A |
| AraC-TAL14 | ✗ | ✓ | ✓ | ✓ | ✓ | ✗ | ✗ | N/A |
| AraC-TAL15 | ✗ | ✓ | ✓ | ✓ | ✓ | ✗ | ✗ | N/A |
| AraC-TAL16 | ✗ | ✓ | ✓ | ✓ | ✓ | ✗ | ✗ | N/A |
| AraC-TAL17 | ✗ | ✓ | ✓ | ✓ | ✓ | ✗ | ✗ | N/A |
| AraC-TAL<sup>+</sup>/OA<sup>-</sup> | N/A | ✓ | N/A | ✓ | ✗ | N/A | N/A | ✓ |

✓: fold-induced GFP expression > 2-fold
✗: fold-induced GFP expression < 2-fold
N/A: not tested

While a sensor that responds to OA and not TAL is useful for engineering OA biosynthesis, the opposite specificity is also of interest for discriminating between the two compounds (e.g. to probe chain elongation in an OA-producing PKS). As a proof of principle, the same random mutagenesis library as above (AraC-TAL1 as parent) was again enriched using the *tetA* dual selection platform, but this time with the inclusion of 2 mM OA as a “decoy” compound during negative selection steps. Following three rounds of selection/counterselection, one interesting variant named AraC-TAL^+^/OA^-^ was isolated and characterized. AraC-TAL^+^/OA^-^ shows slightly lowered background expression compared to AraC-TAL1, a slightly reduced response to TAL, and no response to OA (Figure 2). Further investigation revealed that OA in fact inhibits the ability of TAL to induce AraC-TAL^+^/OA^-^ (addition of 0.5 mM OA to the culture prevents GFP expression in response to TAL), while no such inhibition is seen with AraC-TAL1 (Figure 2C). Addition of OA also further reduces background expression caused by AraC-TAL^+^/OA^-^, to a level near that of wild-type AraC. Interestingly, AraC-TAL^+^/OA^-^ carries four amino acid substitutions compared to parent AraC-TAL1 (Table 1). Analysis of these substitutions, including characterization of AraC-TAL1 variants carrying each single substitution, is provided in the supporting information (Figure S4).

**Figure 2.**
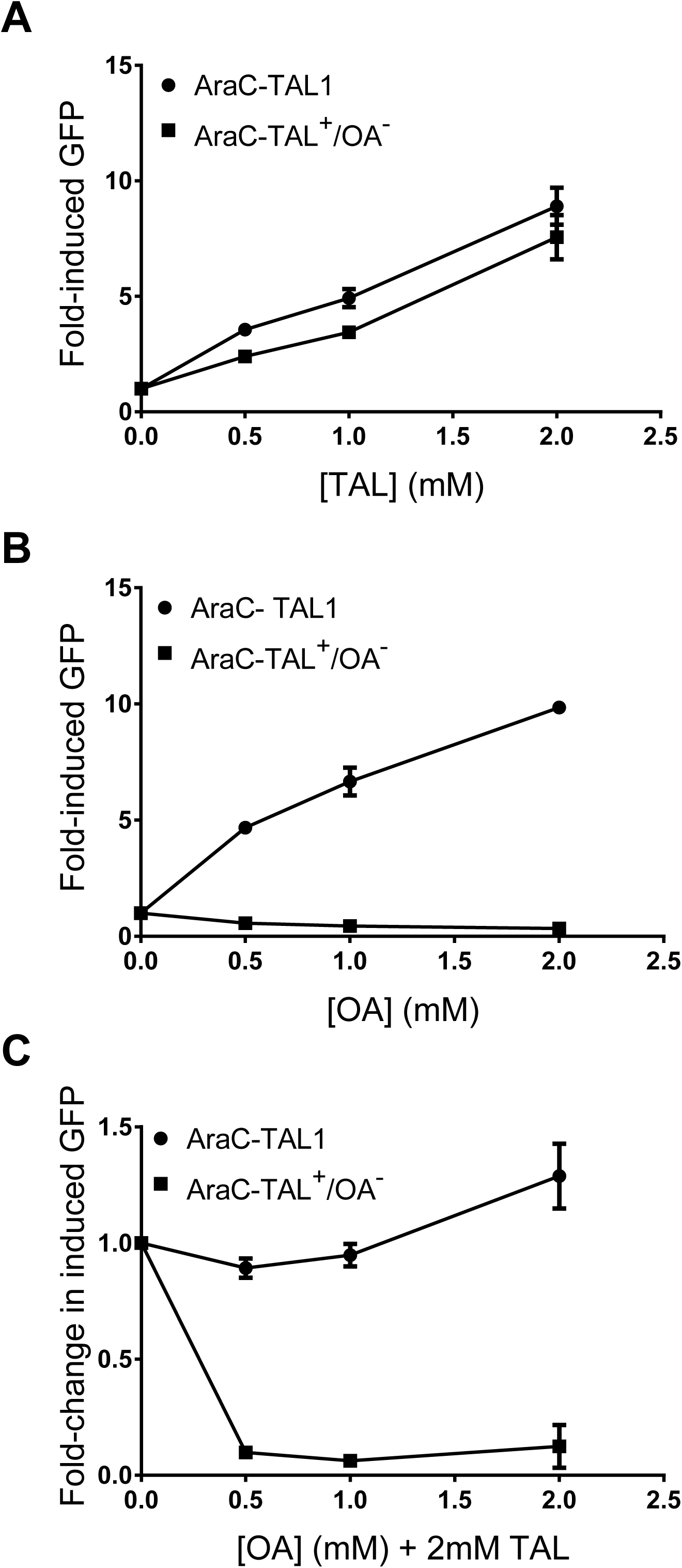
Induced GFP expression of AraC-TAL1 and AraC-TAL^+^/OA^-^ induced with A) triacetic acid lactone (TAL), B) orsellinic acid (OA), and C) OA in presence of 2 mM TAL. Data points are the average of two values, and error bars represent the range.

While OA inhibits gene activation by AraC-TAL^+^/OA^-^, this same variant still shows a strong induced response to 6-methyl-2-hydroxybenzoic acid (differs from OA by one hydroxyl group) and 4- hydroxybenzoic acid (differs from OA by one hydroxyl and one methyl group) (Table 2). Inhibition of AraC- TAL^+^/OA^-^ by OA is reflective of inhibition of wild-type AraC by D-fucose (Doyle et al. 1972), where D-fucose differs from the native inducer L-arabinose by one methyl group. Structural analysis comparing the AraC LBD in complex with L-arabinose vs. D-fucose highlights how relatively subtle differences in ligand binding can result in inhibition of AraC rather than gene activation, which requires major conformational changes (Doyle et al. 1972). Modeling studies to investigate those molecular determinants of ligand binding which correlate with induced gene expression vs. inhibition of gene expression are beyond the scope of the present study, but are anticipated to provide new insights into rational design of AraC-based sensors.

### Design of a Sensor with Enhanced Specificity toward Orsellinic Acid

Among other PKS systems, orcinol synthase from *Rhododendron dauricum* (RdOrs) is a type III PKS that produces orsellinic acid (OA), using acetyl-CoA and malonyl-CoA starter molecules (Taura et al. 2016). OA biosynthesis proceeds through the formation of a tetraketide intermediate, with TAL appearing as a byproduct due to lactonization of the triketide precursor (Taura et al. 2016). The type III PKS 2-pyrone synthase (2-PS) from *Gerbera hybrida* similarly uses acetyl-CoA and malonyl-CoA starter molecules to proceed through a triketide intermediate in the biosynthesis of TAL, with no tetraketide product observed (Austin and Noel 2003). Such control over the length of the final polyketide product is often attributed to the shape and volume of the active site cavity of the type III PKS (Austin and Noel 2003). A biosensor that responds to OA and not TAL could therefore be a useful tool, both for engineering enhanced OA biosynthesis in a recombinant host, and for probing/engineering chain length control and substrate/product specificity in these and similar type III PKS enzymes. We therefore sought to alter the ligand specificity of an OA-responsive AraC-TAL variant.

While using the *tetA* dual selection system with addition of OA as decoy ligand proved effective for isolating the AraC-TAL^+^/OA^-^ variant, this was deemed quite fortuitous: during the end-point screening with the fluorescence assay, only 2 out of 94 clones selected from the enriched library showed response to TAL, while the rest of the population was minimally responsive to TAL and OA. Both TAL-responsive clones were then identified to be identical. In retrospect, considering OA and TAL induce the parent (AraC- TAL1) to similar extents, a single library of random substitution variants was more likely to yield an OA- inhibited variant using this selection method, as compared to a variant that shows little response to OA while retaining strong response to TAL. Prior to further directed evolution, we therefore turned to computational modeling and binding calculations for insights into improving specificity toward OA.

Computational models of AraC-TAL variants (1, 12, 14, 15, and 19) were generated using the Mutator plugin of the IPRO program (Pantazes et al. 2015) by imposing the appropriate amino acid substitutions onto the structure of wild-type AraC (PDB accession: 2ARC (Soisson et al. 1997)), followed by a CHARMM- energy minimization step for clash-free re-packing of the amino acid side chains that lie within 10 Å of the altered residue. Initial docked conformation of OA in AraC and the list of neighboring (within 10Å) pocket residues were obtained by using the *find_contacts* module of OptMAVEn-2.0 program (Chowdhury, Allan and Maranas 2018). Interaction energy scores between the AraC variants and ligands TAL and OA were computed using the CHARMM energy function (Brooks et al. 2009) and used as a proxy for binding affinity. Table S2 lists the CHARMM interaction energy scores (sum of van der Waals forces, electrostatics, and implicit solvation effects modeled as a dielectric continuum), along with the corresponding experimentally measured fold-increases in GFP expression in response to 1 mM inducer (TAL or OA), for these AraC-TAL variants as well as wild-type AraC. The relative ratios of CHARMM interaction energy scores (OA/TAL, also presented in Table S2) correlate well with the ratios of fold-induced GFP expression (OA/TAL) (R^2^ = 0.75), indicating that relative interaction energy scores reasonably capture ligand specificity. Of the variants, AraC-TAL14 demonstrated the strongest response to OA, while still showing low background GFP expression in the absence of TAL (Figures 1B, S1, S2B). AraC-TAL14 was therefore chosen as the starting point for computational design.

OA-specific AraC variants were predicted using the Iterative Protein Redesign and Optimization suite of programs -IPRO (Pantazes et al. 2015); refer to Methods section). IPRO has been previously used to successfully tune substrate and cofactor specificities of several enzymes including bacterial thioestereases (Hernández Lozada et al. 2018, Grisewood et al. 2017) and non-ribosomal peptide synthases (Throckmorton et al. 2019). The top five designs (OA/TAL interaction energy ratios) as identified by IPRO, named AraC-OA1 through AraC-OA5, are listed in Table S3. Note that AraC-OA1, 3, and 4 each has a single amino acid substitution relative to AraC-TAL14, while AraC-OA2 and AraC-OA5 each have two substitutions relative to AraC-TAL14. These five variants were constructed and subsequently characterized experimentally. The experimentally determined specificity for each IPRO-designed variant, here defined as GFP expression response to 1.5 mM OA relative to that in the presence of 3 mM TAL, is given in Table S3 (fold-induced GFP expression for all variants is provided in Table S3 as well). Only AraC- OA1 and AraC-OA4 showed significant response to OA (>2-fold change in fluorescence when induced), and none of these variants showed higher specificity toward OA than AraC-TAL14.

While none of the IPRO-predicted variants tested showed higher specificity than AraC-TAL14, we felt it’s prudent to screen a small library comprising all combinations of the five single or double amino acid substitutions returned by IPRO (those in AraC-OA1 through AraC-OA5), with AraC-TAL14 as parent (32 combinations). Using our P_BAD_-*gfp* reporter construct, 450 clones were screened by microtiter plate- based fluorescence assay, in the presence vs absence of 3 mM TAL or 0.5 mM OA. Screening was carried out with the intent to isolate variants with low TAL response, high OA response, and low background GFP expression. Most variants screened showed either no response to OA or TAL, or showed high background GFP expression. However, four somewhat promising clones were selected for further characterization. Sequencing revealed that these four clones were identical, carrying substitutions P25G and N139G. As shown in Table 3, this variant shows low background expression and low GFP expression response to 3 mM TAL, but also a reduced response to 1.5 mM OA, as compared to AraC-TAL14. Substitution of P25G or N139G (individually, in AraC-TAL14) revealed that P25G alone increases OA specificity to a level similar to that of the AraC-TAL14 parent, and also reduces response to TAL (Table 3). AraC-TAL14/P25G was renamed AraC-OA6 and this variant was selected as the parent for engineering further improvements in OA response and specificity via directed evolution.

**Table 3.** OA specificities of variants identified from the combinatorial library comprising all combinations of amino acid substitutions returned by IPRO.

| **AraC-TAL14**<br>**variants** | **Background GFP**<br>**expression** | **Fold-induced GFP**<br>**expression with**<br>**1.5 mM OA (N=3,**<br>**± SD) (A)** | **Fold-induced GFP**<br>**expression with**<br>**3.0 mM TAL (N=3,**<br>**± SD) (B)** | **OA Specificity**<br>**(A/B)** |
| --- | --- | --- | --- | --- |
| AraC-TAL14 | 116 ± 13 | 6.6 ± 0.3 | 10 ± 0.5 | 0.7 |
| AraC-TAL14/<br>P25G/N139G | 118 ± 14 | 3.8 ± 0.4 | 4.6 ± 0.3 | 0.8 |
| AraC-TAL14/<br>P25G (AraC-OA6) | 223 ± 12 | 3.6 ± 0.5 | 3.8 ± 0.3 | 0.9 |
| AraC-TAL14/<br>N139G | 84 ± 11 | 3.1 ± 0.2 | 6.9 ± 0.3 | 0.5 |

### Directed Evolution of an OA-Specific AraC Variant

A library of AraC-OA6 variants carrying random substitution in the ligand binding domain was generated was screened with a microtiter plate-based fluorescence assay, using our P_BAD_-*gfp* reporter system. The library was first screened in the presence vs absence of 1.5 mM OA, followed by analysis of response to TAL by selected clones. From this first round of screening (513 transformants were screened and four unique clones with high response to OA were further characterized), variant AraC-OA7 showed the highest response to OA (Figure S5A), along with a low response to TAL (6-fold increase in GFP fluorescence in the presence of 3 mM TAL, as compared to 14-fold increase in GFP fluorescence with AraC-TAL14) (Figure S5B), and showed a background expression comparable to wild-type AraC. This resulted in a 10-fold increase in OA specificity as compared to AraC-TAL14, and a 15-fold increase compared to AraC-TAL1 (Figure 3). AraC-OA7 carries three amino acid substitutions relative to its parent (Table 1). Analysis of the individual contributions of each of these three substitutions is presented in Table S4. We next subjected AraC-OA7 to another round of random mutagenesis and screening (513 transformants were screened and six unique clones with high response to OA were further characterized), resulting in AraC-OA8, the variant with the highest specificity towards OA (Figure 3, S6). AraC-OA8 shows a 15-fold increase in OA specificity as compared to AraC-TAL14 (24-fold compared to AraC-TAL1), and retains low background expression (Figure 3). AraC-OA8 carries two amino acid substitutions relative to its parent (Table 1), and 12 total amino acid changes as compared to wild-type AraC. Dose responses of AraC-OA8 to OA is provided in Figure S7. We expect that variants with further improvements in OA specificity and/or sensitivity would be readily discovered through continued rounds of directed evolution.

**Figure 3.**
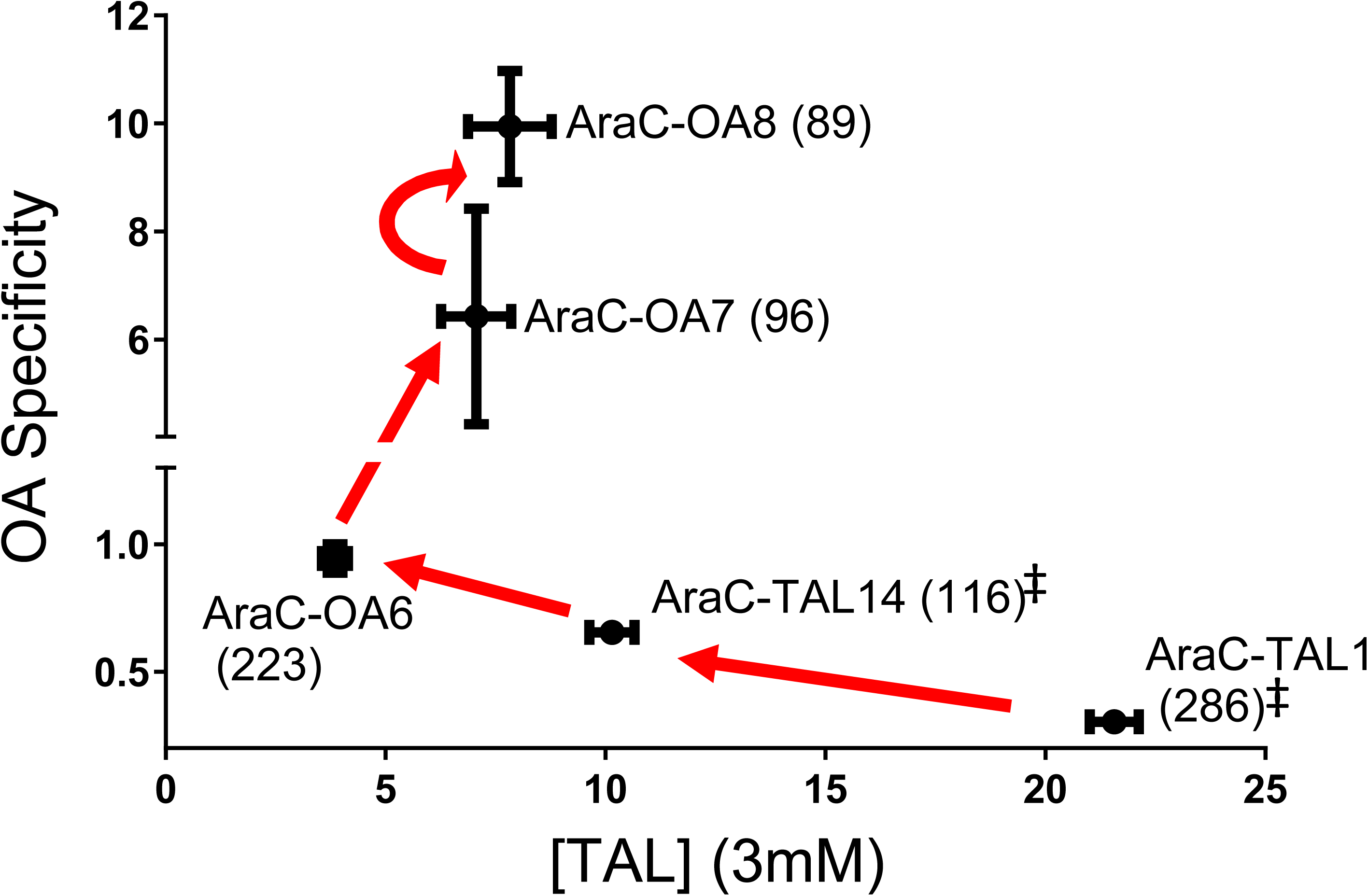
Pathway of evolving OA specificity in AraC-based biosensors. OA specificity is defined as the ratio of fold-induced GFP by 1.5 mM OA vs. fold-induced GFP by 3 mM TAL. Values in parentheses represent background GFP expression normalized by cell density (RFU/OD595) in the absence of inducer. Background values marked with ‡ are average of two values. All other data points are the average of three values and error bars represent standard deviation.

## Discussion

While engineering allosteric regulatory proteins to respond to non-native ligands for biosensor applications is relatively straightforward, attaining significant (or desired) sensitivity and specificity is less clear-cut, requiring more challenging optimization. In this work, we improved upon features of AraC-TAL1, an AraC variant that responds to both TAL and OA, resulting in new variants with greater sensitivity and specificity towards TAL or OA. Table 1 provides the amino acid substitutions found in all AraC variants described in this work. Four images in Figure S8 (A thru D) show the locations of these residues in the AraC crystal structure, in relation to the ligand binding pocket. Understanding how amino acid substitutions affect gene expression by AraC is complicated by the fact that ligand binding must also induce significant conformational changes, including changes in interactions between the ligand binding and DNA binding domains, before gene expression is activated (Soisson et al. 1997). Below we merely highlight salient features of some substitutions that were identified.

We first isolated six AraC-TAL1 variants with reduced background expression levels and improved sensitivity to TAL, from a random substitution library of the AraC-TAL1 ligand binding domain. All variants isolated in this round bear single amino acid substitutions, and display different levels of background expression, sensitivity to TAL (Figure S1), and substrate specificities (Figure S2). AraC-TAL12 shows the highest response to TAL amongst all existing TAL sensors (Figure S1), and varies from AraC-TAL1 only by substitution P11L, located in the AraC N-terminal arm (comprised of residues 8-20). This same substitution reduces background expression compared to AraC-TAL1. The effect of P11L is intriguing in that most substitutions in the N-terminal arm lead to high levels of leaky expression (Berrondo, Gray and Schleif 2010, Dirla, Chien and Schleif 2009). Perhaps in the context of AraC-TAL1, P11L helps to stabilize the repressing conformation in the absence of TAL (strengthens interaction with the DNA binding domain), while also enhancing TAL binding (Saviola, Seabold and Schleif 1998, Soisson et al. 1997). AraC-TAL14 (L72V) and AraC-TAL15 (G135W) show the lowest background expression levels, comparable to that of wild-type AraC (Figure S1). Position 72 is in proximity to AraC ligand binding pocket (Seedorff and Schleif 2011) (Figure S8B); this conservative substitution may help stabilize the AraC repressing conformation, without significantly affecting inducibility by TAL. Meanwhile substitution G135W lies in the AraC dimerization interface (Figure S8D) (Seedorff and Schleif 2011). Substitutions in the dimerization interface have been noted to indirectly enhance AraC repression (Tang and Cirino 2010).

As our TAL-responsive AraC variants also respond to OA, we reasoned that one of these would make a suitable starting point for engineering an OA-specific biosensor (e.g. as opposed to starting with wild-type AraC, which shows no response to OA). To this end, we started with AraC-TAL14 as the parent, and identified variants with improved specificity towards OA. Variant AraC-OA6, with substitution P25G, shows reduced gene expression response to both TAL and OA, but also higher specificity toward OA (Figure 3). Residue 25 resides in the first β-strand of the ligand binding pocket (Figure S8B). Substitution P25S in wild-type AraC was previously reported to reduce response to arabinose by 4-fold, possibly due to weakened interactions between the N-terminal arm and the DNA binding domain in the activating conformation (Reed and Schleif 1999). Variant AraC-OA7 bears three additional amino acid substitutions (I36F, P128S, and A140V). Results from characterizing each single back-substitution within AraC-OA7 are provided in Table S4. Of most significance, while I36F alone enhances response towards OA, all three new substitutions are necessary to achieve the highest specificity and lowest background expression. A final round of evolution yielded variant AraC-OA8, with two additional substitutions. In total, AraC-OA8 contains twelve amino acid substitutions compared to wild-type AraC, shows 24-fold improved specificity toward OA over TAL relative to AraC-TAL1. This variant retains repression in the absence of inducer to the same extent as wild-type AraC, and shows a 6-fold induced GFP expression response to 125 μM OA. This new biosensor/reporter should prove useful for engineering OA biosynthesis in *E. coli*.

