## Supplementary materials for "Engineering sensitivity and specificity of AraC-based biosensors responsive to triacetic acid lactone and orsellinic acid"

**Affiliations:**

### Table of Contents

**Plasmid map and sequence** of pPCC1322

**Table S1**. Characterization of variant AraC-TAL18 carrying substitutions from AraC-TAL12 (P11L) and AraC-TAL17 (Q54R)

**Table S2**. Relative CHARMM interaction energy scores and experimentally measured specificity ratios of AraC-TAL1 variants towards OA and TAL

**Table S3**. Relative interaction scores and specificity ratios for computationally-predicted OA-preferring variants: CHARMM calculations vs. experimental results

**Table S4**. Fold-induced GFP expression values and OA specificities of AraC-TAL14 variants bearing single amino acid substitutions identified in AraC-OA7

**Figure S1**. Dose response curves of AraC-TAL1 variants with varying TAL concentrations

**Figure S2**. Dose response curves of AraC-TAL1 variants towards various compounds with similar structures to TAL

**Figure S3**. Dose response curves of AraC-$TAL^{+}$/$OA^{-}$ variant in comparison with AraC-TAL1 for OA and OA analogues

**Figure S4**. Dose response curves of AraC-TAL1 variants bearing single amino acid substitutions identified in AraC-$TAL^{+}$/$OA^{-}$ variant in comparison with AraC-TAL1 for TAL and OA

**Figure S5**. Fold-induced GFP values of the four unique variants isolated from random mutagenesis of AraC-OA6 towards TAL and OA

**Figure S6**. Fold-induced GFP values of the six unique variants isolated from random mutagenesis of AraC-OA7 towards TAL and OA

**Figure S7**. Fold-induced GFP values of AraC-OA8, in comparison with ArC-TAL14, when induced with OA

**Figure S8**. Relative positions of amino acid substitutions found in AraC-TAL1 variants identified in this work

Plasmid map and full sequence of pPCC1322. Sequence in red indicates AraC-TAL1 coding sequence, and sequence in green indicates GPF coding sequence.

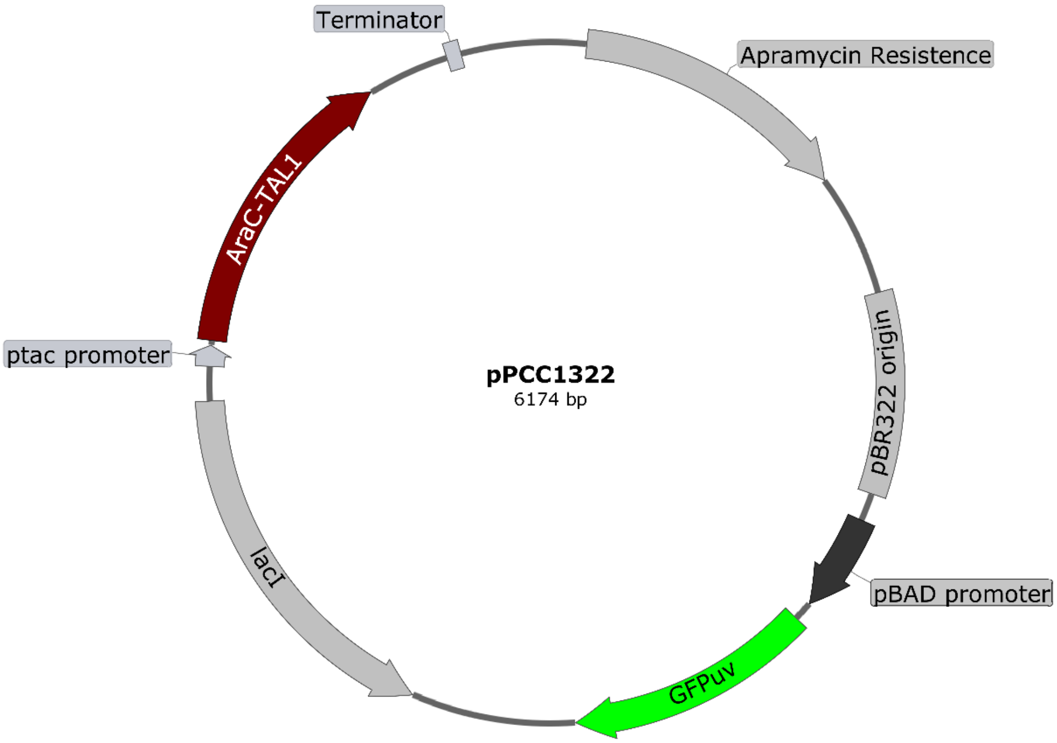


ccggcctttgaatgggttcatgtgcagctccatcagcaaaaggggatgataagtttatcaccaccgactatttgcaacagtgccgttgatcgtgctatga
tcgactgatgtcatcagcggtggagtgcaatgtcgtgcaatacgaatggcgaaaagccgagctcatcggtcagcttctcaaccttggggttaccccg
gcggtgtgctgctggtccacagctccttccgtagcgtccggcccctcgaagatgggccacttggactgatcgaggccctgcgtgctgcgctgggtccgg
gagggacgctcgtcatgccctcgtggtcaggtctggacgacgagccgttcgatcctgccacgtcgcccgttacaccggaccttggagttgtctctgaca
cattctggcgcctgccaaatgtaaagcgcagcgcccatccatttgcctttgcggcagcggggccacaggcagagcagatcatctctgatccattgcccc
tgccacctcactcgcctgcaagcccggtcgcccgtgtccatgaactcgatgggcaggtacttctcctcggcgtgggacacgatgccaacacgacgctgc
atcttgccgagttgatggcaaaggttccctatggggtgccgagacactgcaccattcttcaggatggcaagttggtacgcgtcgattatctcgagaatg
accactgctgtgagcgctttgccttggcggacaggtggctcaaggagaagagccttcagaaggaaggtccagtcggtcatgcctttgctcggttgatcc
gctcccgcgacattgtggcgacagccctgggtcaactgggccgagatccgttgatcttcctgcatccgccagagggcgggatgcgaagaatgcgatgc
cgctcgccagtcgattggctgagctcatgagcggagaacgagatgacgttggaggggcaaggtcgcgctgattgctggggcaacacatcattgcagc
actggggccagatggtaagccctcccgtatcgtagttatctacacgacggggagtcaggcaactatggatgaacgaaatagacagatcgctgagata
ggtgcctcactgattaagcattggtaactgtcagaccaagtttactcatatatactttagattgatttaaaacttcatttttaatttaaaaggatctaggtg
aagatcctttttgataatctcatgaccaaaatcccttaacgtgagttttcgttccactgagcgtcagaccccgtagaaaagatcaaaggatcttcttgag
atcctttttttctgcgcgtaatctgctgcttgcaaacaaaaaaaccaccgctaccagcggtggtttgtttgccggatcaagagctaccaactctttttccg
aaggtaactggcttcagcagagcgcagataccaaatactgtccttctagtgtagccgtagttaggccaccacttcaagaactctgtagcaccgcctaca
tacctcgctctgctaatcctgttaccagtggctgctgccagtggcgataagtcgtgtcttaccgggttggactcaagacgatagttaccggataaggcgc
agcggtcgggctgaacggggggttcgtgcacacagcccagcttggagcgaacgacctacaccgaactgagatacctacagcgtgagctatgagaaa
gcgccacgcttcccgaagggagaaaggcggacaggtatccggtaagcggcagggtcggaacaggagagcgcacgagggagcttccagggggaaa
cgcctggtatctttatagtcctgtcgggtttcgccacctctgacttgagcgtcgatttttgtgatgctcgtcaggggggcggagcctatggaaaaacgcc
agcaacgcggcctttttacggttcctggccttttgctggccttttgctcacttttcatactcccgccattcagagaagaaaccaattgtccatattgcatc
agacattgccgtcactgcgtcttttactggctcttctcgctaaccaaaccggtaaccccgcttattaaaagcattctgtaacaaagcgggaccaaagcc
atgacaaaaacgcgtaacaaaagtgtctataatcacggcagaaaagtccacattgattatttgcacggcgtcacactttgctatgccatagcatttttta
tccataagattagcggatcctacctgacgctttttatcgcaactctctactgtttctccatacccgtttttttgggctagaaataattttgtttaactttaaga

aggagatatacatatggctagcaaaggagaagaacttttcactggagttgtcccaattcttgttgaattagatggtgatgttaatgggcacaaattttct
gtcagtggagagggtgaaggtgatgctacatacggaaagcttacccttaaatttatttgcactactggaaaactacctgttccatggccaacacttgtc
actactttctcttatggtgttcaatgcttttcccgttatccggatcatatgaaacggcatgactttttcaagagtgccatgcccgaaggttatgtacagga
acgcactatatctttcaaagatgacgggaactacaagacgcgtgctgaagtcaagtttgaaggtgatacccttgttaatcgtatcgagttaaaaggtat
tgattttaaagaagatggaaacattctcggacacaaactcgagtacaactataactcacacaatgtatacatcacggcagacaaacaaaagaatgga
atcaaagctaacttcaaaattcgccacaacattgaagatggatccgttcaactagcagaccattatcaacaaaatactccaattggcgatggccctgtc
cttttaccagacaaccattacctgtcgacacaatctgccctttcgaaagatcccaacgaaaagcgtgaccacatggtccttcttgagtttgtaactgctg
ctgggattacacatggcatggatgagctctacaaataatgaattcgagctcggtaccgggccccccctcgaggtcgacggtatcgataagcttgatatc
gaattcctgcagcccgggggatccactagttctagagcggccgccaccgcggtggagctccagcttttgttccctttagtgagggttaattgcgcaaac
acagaaaaaagcccgcacctgacagtgcgggcttttttttttgcaactgttgggaagggcgatcggtgcgggcctcttcgctattacgccagctggcga
aagggggatgtgctgcaaggcgattaagttgggtaacgccagggttttcccagtcacgacgttgtaaaacgacggccagtgaatccgtaatcatggtc
atagctgtttcctgtgtgaaattgttatccgctcacaattccacacaacatacgagccggaagcataaagtgtaaagcctggggtgcctaatgagtgag
ctaacttacattaattgcgttgcgctcactgcccgctttccagtcgggaaacctgtcgtgccagctgcattaatgaatcggccaacgcgcggggagagg
cggtttgcgtattgggcgccagggtggtttttcttttcaccagtgagacgggcaacagctgattgcccttcaccgcctggccctgagagagttgcagcaa
gcggtccacgctggtttgccccagcaggcgaaaatcctgtttgatggtggttaacggcgggatataacatgagctgtcttcggtatcgtcgtatcccact
accgagatatccgcaccaacgcgcagcccggactcggtaatggcgcgcattgcgcccagcgccatctgatcgttggcaaccagcatcgcagtgggaa
cgatgccctcattcagcatttgcatggtttgttgaaaaccggacatggcactccagtcgccttcccgttccgctatcggctgaatttgattgcgagtgaga
tatttatgccagccagccagacgcagacgcgccgagacagaacttaatgggcccgctaacagcgcgatttgctggtgacccaatgcgaccagatgct
ccacgcccagtcgcgtaccgtcttcatgggagaaaataatactgttgatgggtgtctggtcagagacatcaagaaataacgccggaacattagtgcag
gcagcttccacagcaatggcatcctggtcatccagcggatagttaatgatcagcccactgacgcgttgcgcgagaagattgtgcaccgccgctttaca
ggcttcgacgccgcttcgttctaccatcgacaccaccacgctggcacccagttgatcggcgcgagatttaatcgccgcgacaatttgcgacggcgcgtg
cagggccagactggaggtggcaacgccaatcagcaacgactgtttgcccgccagttgttgtgccacgcggttgggaatgtaattcagctccgccatcg
ccgcttccactttttcccgcgttttcgcagaaacgtggctggcctggttcaccacgcgggaaacggtctgataagagacaccggcatactctgcgacat
cgtataacgttactggtttcacattcaccaccctgaattgactctcttccgggcgctatcatgccataccgcgaaaggttttgcaccattccatgggaatt
cgctagcgcggccgcgagctgttgacaattaatcatcggctcgtataatgtgtggaattgtgagcggataacaatttcacacaggagatacctaggat
ggctgaagcgcaaaatgatgtgctgctgccgggatactcgtttaacgcccatctggtggcgggtttaatcccgattgaggccaacggttatctcgatttt
tttatcgaccgaccgctgggaatgaaaggttatattctcaatctcaccattcgcggtcaggggggtggtgaaaaatcagggacgagaattcgtctgccg
accgggtgatattttgctgttcccgccaggagagattggccacttgggtcgtcatccggaggctcgcgaatggtatcgccagtgggtttactttcgtccg
cgcgcctactggcatgaatggcttaactggccgtcaatatttgccaatacgggtttctttcgcccggatgaagcgcaccagccgcatttcagcgacctgt
ttgggcaaatcattaacgccgggcaaggggaagggcgctattcggagctgctggcgataaatctgcttgagcaattgttactgcggcgcatggaagcg
attaacgagtcgctccatccaccgatggataatcgggtacgcgaagcttgtcagtacatcagcgatcacctggcagacagcaattttgatatcgccagc
gtcgcacagcatgtttgcttgtcgccgtcgcgtctgtcacatcttttccgccagcagttagggattagcgtcttaagctggcgcgaggaccaacgcatta
gtcaggcgaagctgcttttgagcactacccggatgcctatcgccaccgtcggtcgcaatgttggttttgacgatcaactctatttctcgcgagtatttaaa
aaatgcaccggggccagcccgagcgagtttcgtgccggttgtgaagaaaaagtgaatgatgtagccgtcaagttgtcataattggtaacgaatcaga
caattgacggcttgacggagtagcatagggtttgcagaatccctgcggtaccagatctgtcgactacaaggacgatgacgacaagtgaagatcgatct
ctcgatcgagtgagagaagacttgcatgcctgcaggtcgactctagaggatccccgtactatcaacaggttgaactgcggatcttgcggccagctttat
gcttgtaaaccgttttgtgaaaaaatttttaaaataaaaaaggggacctctagggtccccaattaattagtaatataatctattaaaggtcattcaaaa
ggtcatccaccggatcaattcccctgctcgcgcaggctgggtgccaagctctcgggtaacatcaaggcccgatccttggagcccttgccctcccgcacg
atgatcgtgccgtgatcgaaatccagatccttgacccgcagttgcaaaccctcactgatccggctcacggtaactgatgccgtatttgcagtaccagcg
tacggcccacagaatgatgtcacgctgaaaatg

**Table S1**. Characterization of variant AraC-TAL18 carrying substitutions from AraC-TAL12 (P11L) and AraC-TAL17 (Q54R)

| | **Background GFP expression (Fluorescence/OD 595 nm)** | **Average fold-induced GFP values (N=2)** | | | | | | |
|---|---|---|---|---|---|---|---|---|
| | | 0.125 mM TAL | 0.25 mM TAL | 0.375 mM TAL | 0.5 mM TAL | 1 mM TAL | 2 mM TAL | 4 mM TAL |
| WT-AraC | 89 | 0.8 | 0.9 | 0.8 | 0.8 | 0.8 | 0.8 | 1.4 |
| AraC-TAL1 | 286 | 1.2 | 1.7 | 2.4 | 3.1 | 5.7 | 15 | 56 |
| AraC-TAL1/ P11L (AraC-TAL12) | 176 | 1.5 | 2.4 | 4.1 | 6.8 | 13 | 49 | 128 |
| AraC-TAL1/ Q54R (AraC-TAL17) | 177 | 1.4 | 2.4 | 2.7 | 3.0 | 5.8 | 11 | 52 |
| AraC-TAL1/ P11L/Q54R (AraC-TAL18) | 162 | 1.2 | 1.9 | 3.0 | 4.2 | 9.3 | 26 | 118 |

**Table S2**. Relative CHARMM interaction energy scores and experimentally measured specificity ratios of AraC-TAL1 variants towards OA and TAL.

| Variant | CHARMM interaction energy score with OA | Fold-increase in GFP with OA (1 mM) (A) | CHARMM interaction energy score with TAL | Fold - increase in GFP with TAL (1 mM) (B) | *In silico* relative interaction score OA/ TAL | OA specificity (OA/ TAL, 1 mM each) (A/B) |
|---|---|---|---|---|---|---|
| Wild-type | -109.3 | 1.0 | -99.2 | 0.9 | 1.1 | 1.1 |
| AraC-TAL1 | -121.8 | 8.5 | -116.6 | 5.7 | 1.0 | 1.5 |
| AraC-TAL12 | -121.7 | 5.6 | -124.4 | 13.5 | 1.0 | 0.3 |
| AraC-TAL14 | -123.8 | 9.9 | -113.2 | 5.2 | 1.1 | 1.9 |
| AraC-TAL15 | -116.4 | 6.4 | -102.3 | 2.3 | 1.1 | 2.8 |
| AraC-$TAL^{+}/OA^{-}$ | -98.1 | 0.4 | -110.8 | 3.7 | 0.9 | 0.1 |

**Table S3**. Relative interaction scores and specificity ratios for computationally-predicted OA-preferring variants: CHARMM calculations vs. experimental results

| Variant | Amino acid substitutions in AraC-TAL14 | CHARMM interaction energy score with OA | CHARMM interaction energy score with TAL | *In silico* relative interaction score OA/ TAL | Fold increase in GFP with OA (1.5 mM) (N=3, ±SD) | OA Specificity from experiment [OA] (1.5 mM)/ [TAL] (3 mM) |
|---|---|---|---|---|---|---|
| AraC-TAL14 | | -123.8 | -113.2 | 1.1 | 6.6±0.3 | 0.7 |
| AraC-OA1 | V8D | -185.8 | -91.4 | 2.0 | 4.0±0.9 | 0.1 |
| AraC-OA2 | I24D,P25G | -150 | -100.3 | 1.5 | 0.9±0.1 | 0.7 |
| AraC-OA3 | V72D | -140 | -102.9 | 1.4 | 0.9±0.1 | 0.8 |
| AraC-OA4 | L82M | -183.2 | -85.8 | 2.1 | 2.3±0.1 | 0.4 |
| AraC-OA5 | G135L,N139G | -173.4 | -93.4 | 1.9 | 0.8±0.1 | 0.7 |

**Table S4**. Fold-induced GFP expression values and OA specificities of AraC-TAL14 variants bearing single amino acid substitutions identified in AraC-OA7

| | **Background GFP** | **Fold-induced GFP with 1.5 mM OA (N=3, ±SD) (A)** | **Fold-induced GFP with 3.0 mM TAL (N=3, ±SD) (B)** | **OA Specificity (A/B)** |
|---|---|---|---|---|
| AraC-TAL14 | 116 ± 3 | 6.6 ± 0.3 | 10 ± 0.5 | 0.7 |
| AraC-TAL14/P25G (AraC-OA6) | 223 ± 12 | 3.6 ± 0.5 | 3.8 ± 0.3 | 0.9 |
| AraC-TAL14/P25G /I36F/P128S/A140V (AraC-OA7) | 96 ± 4 | 45 ± 14 | 7.1 ± 0.8 | 6.4 |
| AraC-TAL14/P25G/ I36F | 160 ± 14 | 53 ± 4 | 16 ± 2 | 3.4 |
| AraC-TAL14/P25G/ P128S | 268 ± 52 | 5.6 ± 0.1 | 7.4 ± 1.8 | 0.8 |
| AraC-TAL14/P25G/ A140V | 123 ± 20 | 2.4 ± 0 | 3.4 ± 0.4 | 0.7 |
| AraC-TAL14/P25G/ I36F/P128S | 204 ± 30 | 43 ± 2 | 15 ± 2 | 2.9 |
| AraC-TAL14/P25G/ P128S/A140V | 134 ± 10 | 2.4 ± 0 | 3.9 ± 0.2 | 0.6 |
| AraC-TAL14/P25G/ I36F/ A140V | 102 ± 4 | 38 ± 1 | 7.9 ± 1.1 | 4.9 |

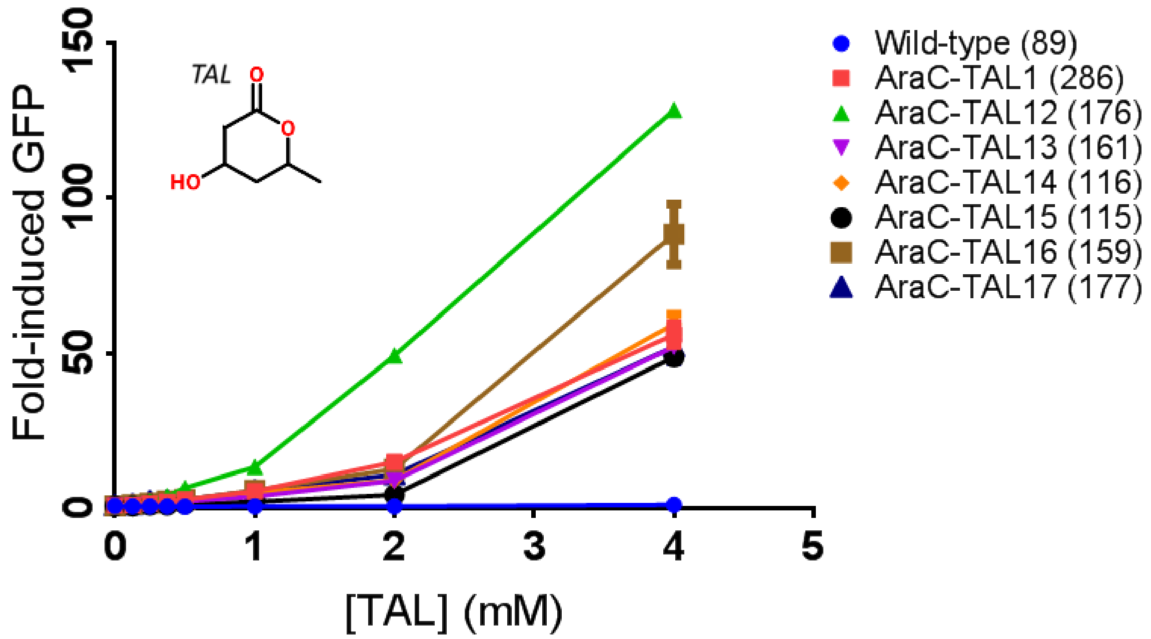


**Figure S1**. Dose response curves of AraC-TAL1 variants towards varying TAL concentrations. Data points are the average of two values, and error bars represent the range. Background GFP expression (GFP expression without TAL) is indicated in parentheses.

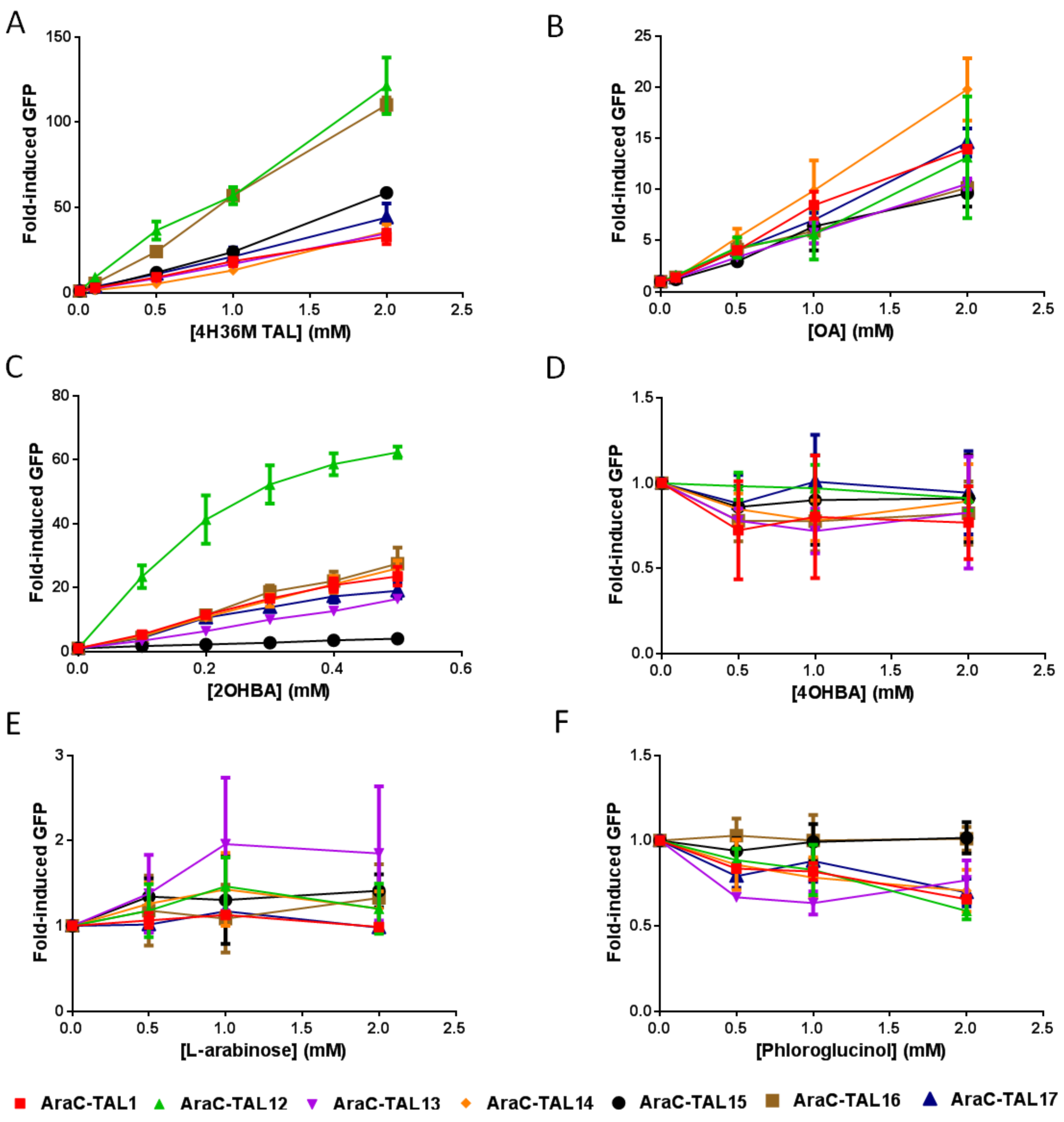


**Figure S2**. Dose response curves of AraC-TAL1 variants towards **A)** 4-Hydroxy-3,6-dimethyl-2-pyrone (4H36M TAL), **B)** Orsellinic acid (OA), **C)** 2-Hydroxybenzoic acid (2OHBA or salicylic acid), **D)** 4-Hydroxybenzoic acid (4OHBA), **E)** L-arabinose, and **F)** Phloroglucinol. Data points are the average of two values, and error bars represent the range.

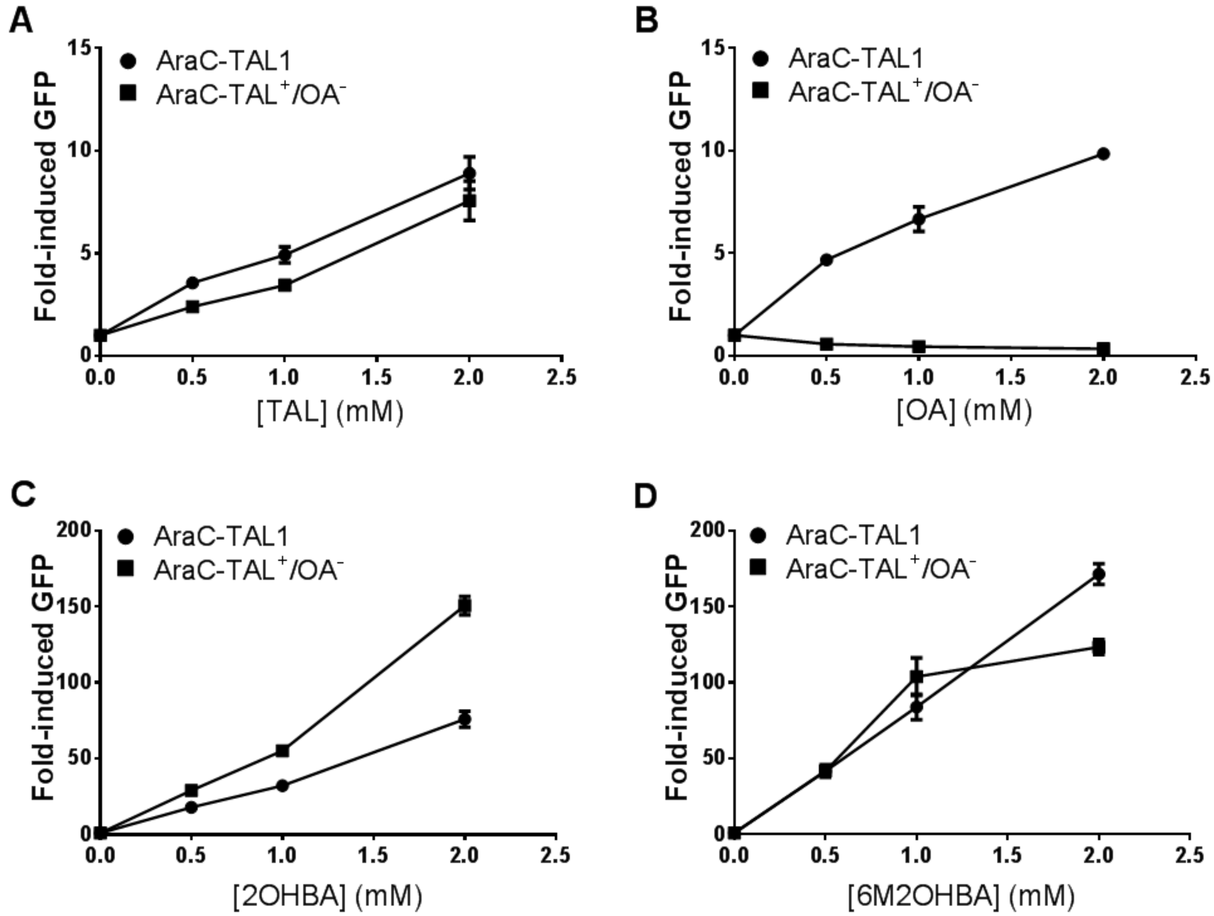


**Figure S3**. Dose response curves of AraC-TAL$^{+}$/OA$^{-}$ variant in comparison with AraC-TAL1 for **A)** Triacetic acid lactone (TAL), **B)** Orsellinic acid (OA) **C)** 2-Hydroxybenzoic acid (2OHBA or salicylic acid), **D)** 6-Methyl-2-hydroxybenzoic acid (6M2OHBA or 6-methyl salicylic acid). Data points are the average of two values, and error bars represent the range.

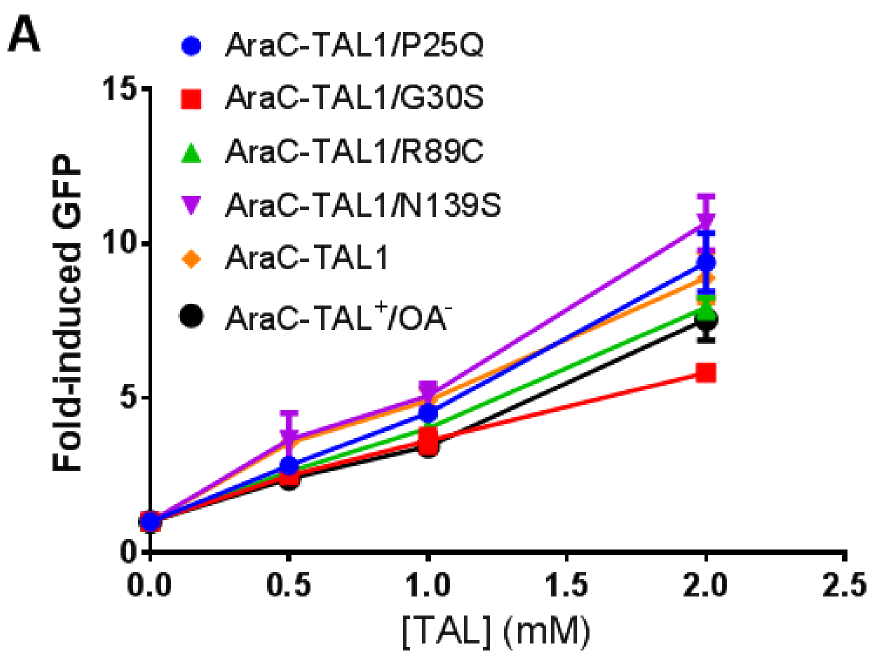


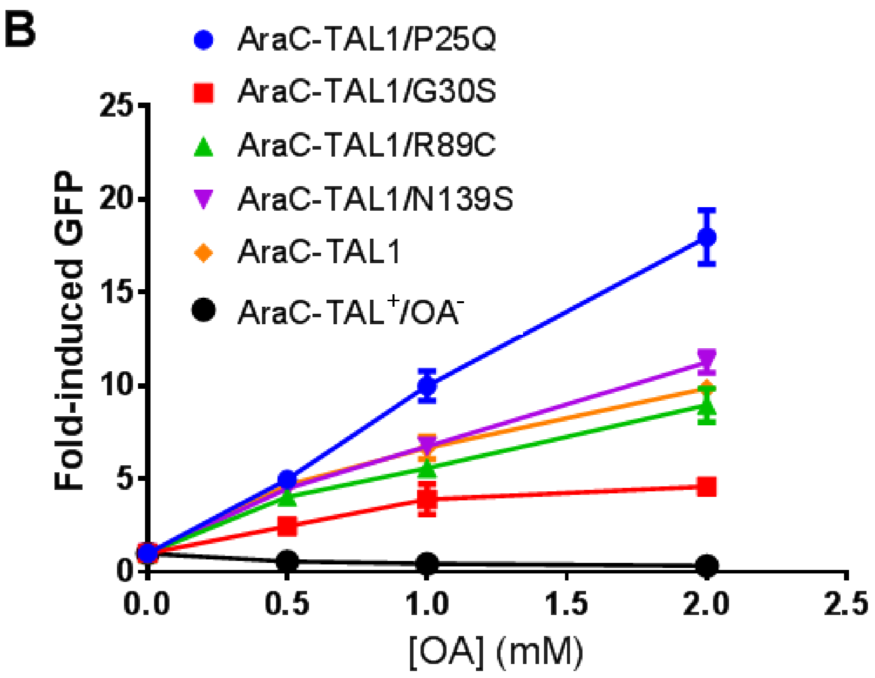


**Figure S4**. Dose response curves of AraC-TAL1 variants bearing single amino acid substitutions identified in AraC-TAL$^{+}$/OA$^{-}$ variant in comparison with AraC-TAL1 for **A)** Triacetic acid lactone (TAL), **B)** Orsellinic acid (OA). Data points are the average of two values, and error bars represent the range.

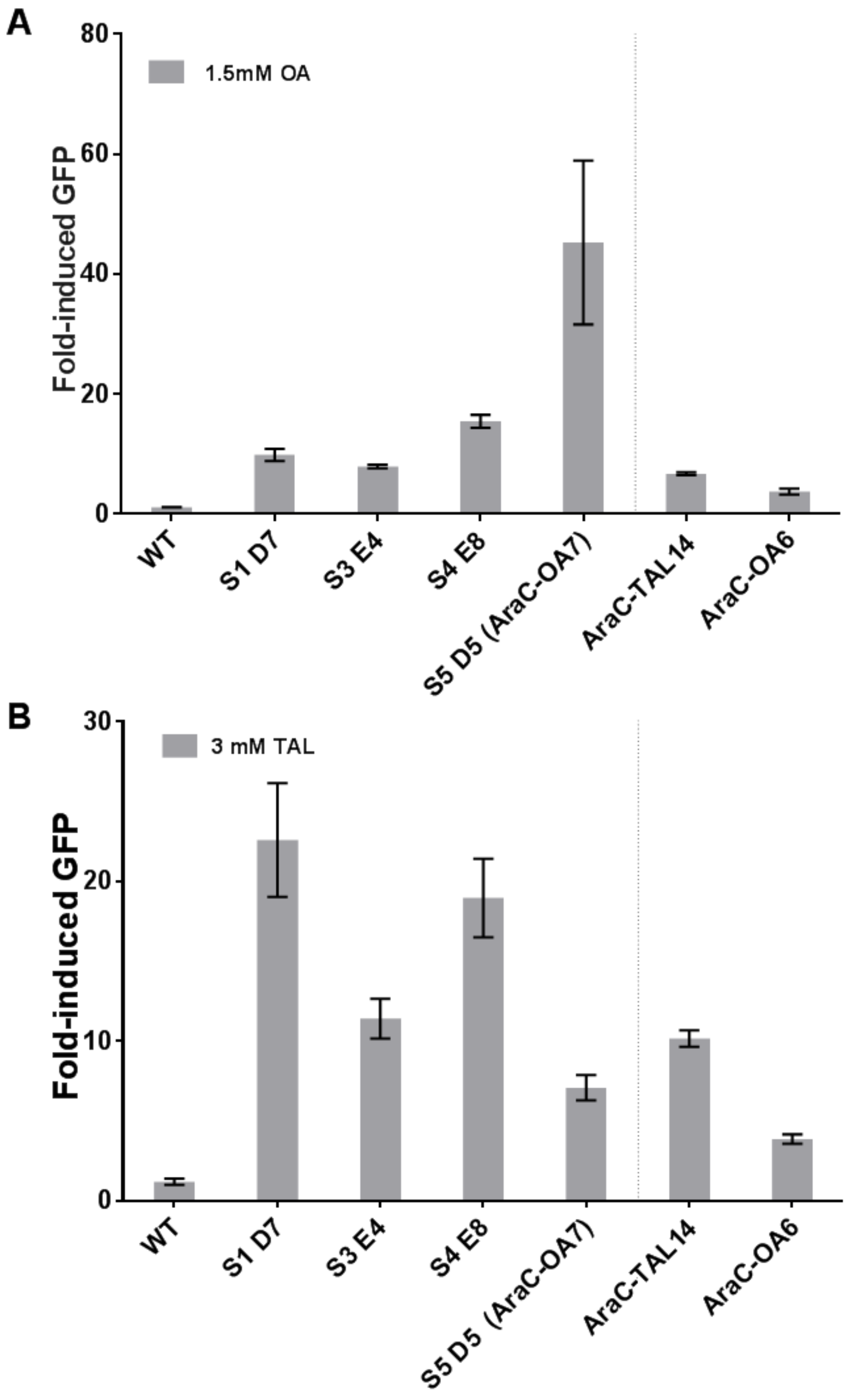


**Figure S5**. Fold-induced GFP values of the four unique variants isolated from random mutagenesis of AraC-OA6 when induced with **A)** OA and **B)** TAL.

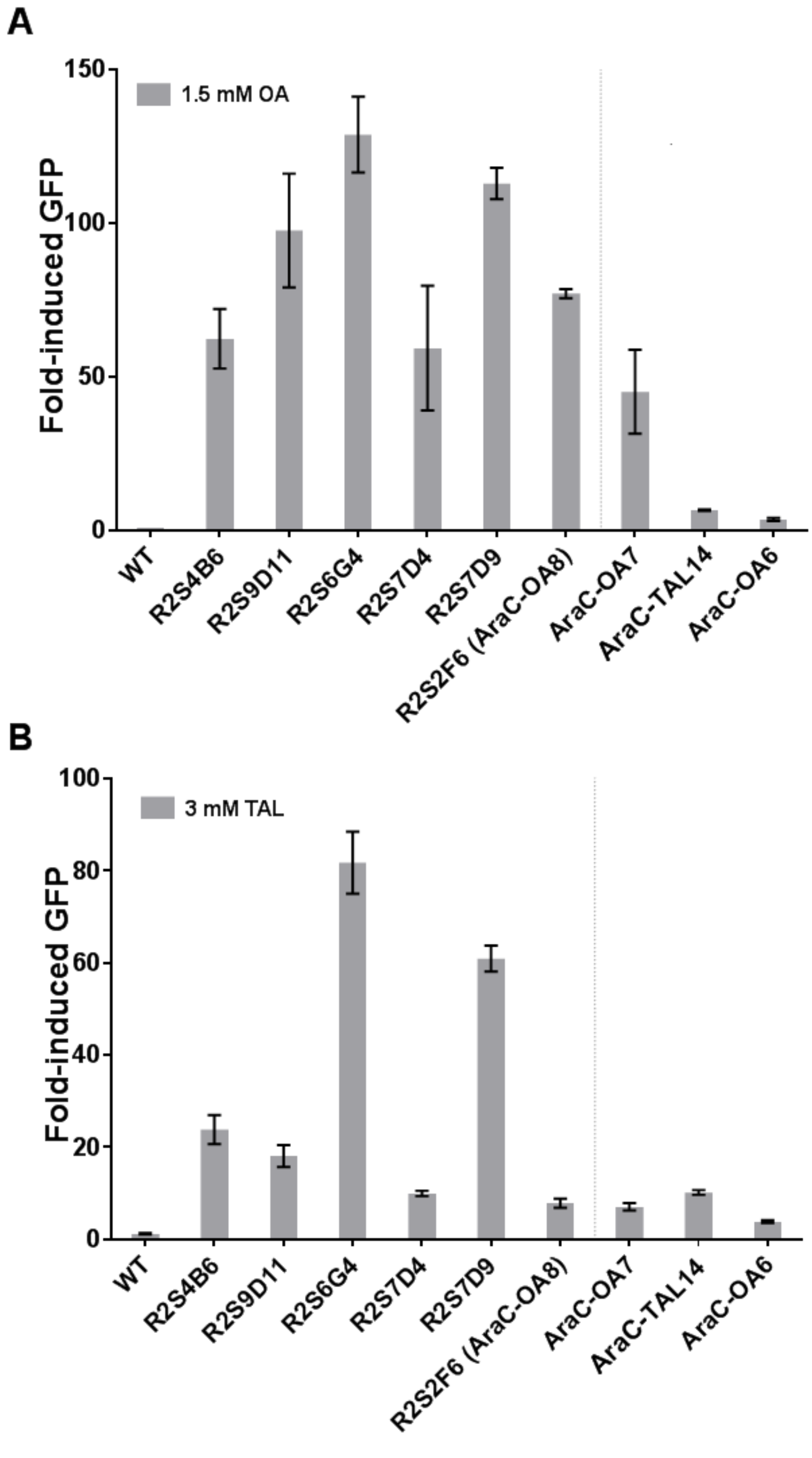


**Figure S6**. Fold-induced GFP values of the six unique variants isolated from random mutagenesis of AraC-OA7 when induced with **A)** OA and **B)** TAL.

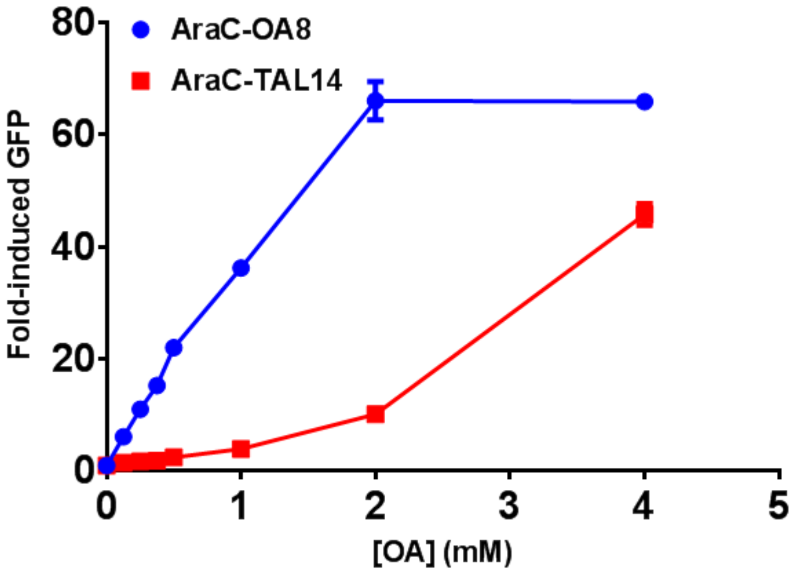


**Figure S7**. Fold-induced GFP values of AraC-OA8, in comparison with ArC-TAL14, when induced with OA.

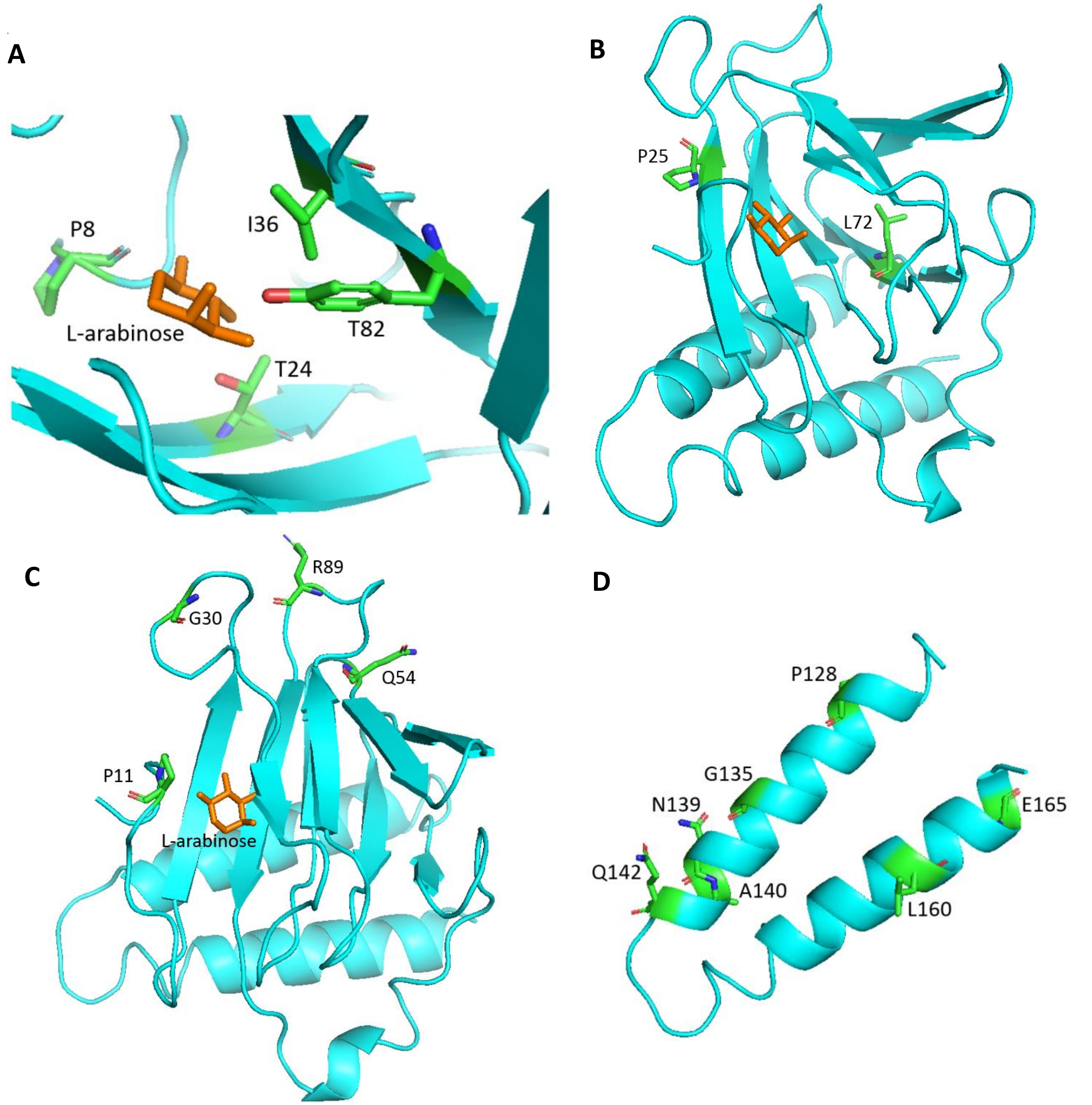


**Figure S8**. Relative positions of amino acid substitutions found in AraC-TAL12 to AraC-TAL17, AraC-$TAL^{+}/OA^{-}$, and AraC-OA1 to AraC-OA8, as summerized in Table 1. **A)** Four amino acid substitutions (V8D, I24D, I36F, and L82M) found in the binding pocket of AraC, based on the crystal strcuture of wild-type AraC (2ARC). The residues in the image are showing the original residues in wild-type AraC. In AraC-TAL1, these residues are V8, I24, I36, and L82. **B)** Two amino acid substitutions that are in proximity to the binding pocket: L72 (3.7 Å), P25 (2.8 Å). **C)** Four amino acid substitutions that were identified to be in the ligand binding domain but not on the dimerization domain: P11 (4.4 Å), G30 (9.9 Å), Q54 (8.3 Å), R89 (10.9 Å). **D)** Seven substitutions in the dimerization domain. Distances in parenthesis indicate the distance from the residue to the binding pocket based wild-type AraC structure (2ARC).
